# Geometric Effects Position Renal Vesicles During Kidney Development

**DOI:** 10.1101/2022.08.30.505859

**Authors:** Malte Mederacke, Lisa Conrad, Nikolaos Doumpas, Roman Vetter, Dagmar Iber

## Abstract

During kidney development, reciprocal signalling between the epithelium and the mesenchyme coordinates nephrogenesis with branching morphogenesis of the collecting ducts. The mechanism that positions the renal vesicles, and thus the nephrons, relative to the branching ureteric buds has remained elusive. By combining computational modelling and experiments, we show that geometric effects concentrate the key regulator, WNT9b, at the junctions between parent and daughter branches where renal vesicles emerge, even when uniformly expressed in the ureteric epithelium. This curvature effect might be a general paradigm to create non-uniform signalling in development.

## Introduction

Developmental processes must be coordinated in space and time to form a functional organ. In the kidney, the nephrons and the collecting ducts develop from different parts of the intermediate mesoderm [11]. Yet, the processes are coordinated such that each nephron connects to a different branch element of the ureteric tree — a design that ensures efficient drainage of the collected fluid. While the key molecular regulators of branching morphogenesis and nephrogenesis have been defined [11, 24], the mechanism that positions the nephrons relative to the ureteric tree has remained elusive. Nephrons develop from renal vesicles (RVs), which in turn emerge from the epithelisation of mesenchymal cell condensations, so-called pretubular aggregates (PTAs) [36]. The PTA/RVs form in the corners between parent branches and newly emerging daughter branches during kidney branching morphogenesis (Fig. 1A).

**Figure 1:**
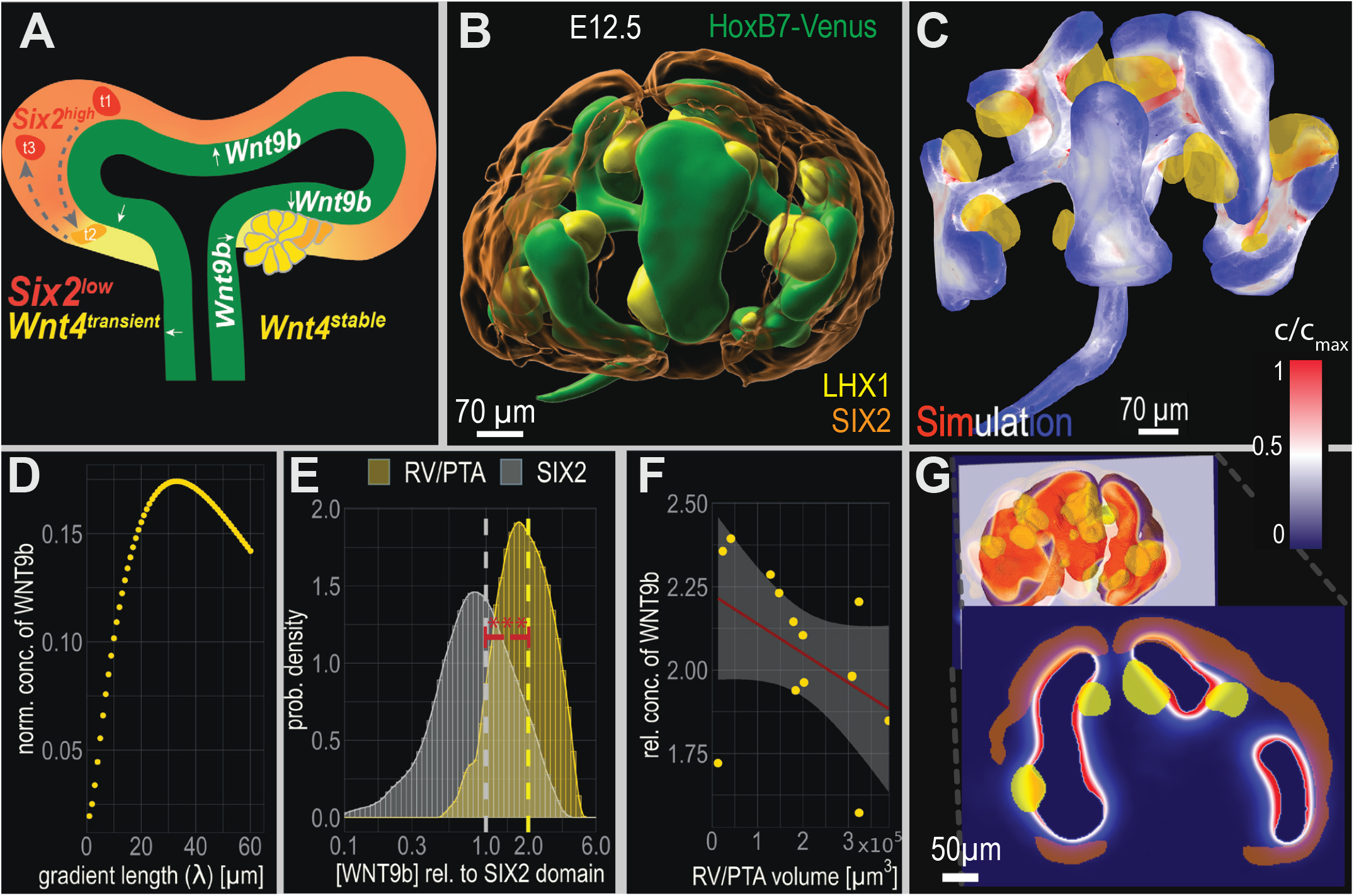
Geometric effects position PTA/RVs in developing kidneys. (A) Regulation of PTA/RV formation. For details, see text. (B) Volumetric light sheet microscopy data of an E12.5 embryonic kidney with surface segmentations for the PTA/RVs (LHX1, yellow), the UB (HoxB7, green) and the CM (SIX2, orange). (C) 3D simulation of the steady-state WNT9b distribution. Uniform secretion of WNT9b from the UB surface (Neumann boundary condition) leads to the highest ligand concentration (red) in the UB corner regions, coinciding with the position of PTA/RVs (yellow). The geometries of the PTA/RVs and the UB were extracted from LSM images. The WNT9b gradient length was set to λ = 30 µm. The colourmap shows the normalised, integrated WNT9b concentration profile along the normal direction of the UB towards the mesenchyme projected back on the surface of the UB. (D) The highest ligand upconcentration in PTA/RVs is achieved for λ ≈ 30 µm. The normalised WNT9b concentration is calculated as the difference between the mean concentration inside the PTA/RVs, 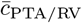 and the mean concentration inside the SIX2 population, 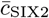, relative to the highest concentration inside the kidney, 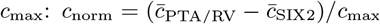. (E) Relative simulated WNT9b concentration inside the PTA/RVs (yellow) and in the SIX2 population (gray). The predicted mean concentration in PTA/RVs (yellow dashed line; 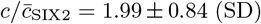) is above the predicted mean concentration in the SIX2 population (grey dashed line; 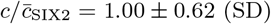) (*p* < 0.001, Welch’s two-sample t-test). (F) The predicted WNT9b concentration inside the PTA/RVs relative to the SIX2 population is higher in smaller PTA/RVs. The red line and gray shade represent a linear fit with its standard error (*R*^2^ = 0.2). (G) A cross-section highlighting the relative location of the CM (orange), PTA/RVs (yellow), and the simulated concentration gradient (blue-red colourmap).

WNT9b is the most upstream regulator of nephrogenesis that has hitherto been identified, and in its absence, no PTA/RVs are observed [4]. *Wnt9b* is expressed largely uniformly in the ureteric epithelium [8], and diffuses to the adjacent metanephric mesenchyme, where it controls PTA/RV formation via canonical WNT signalling [4, 20, 23, 45]. The response of nephron progenitor cells (NPCs) to *β*-catenin, the key effector of canonical WNT signalling, is activity dependent [19, 48]. Low *β*-catenin activity supports NPC maintenance and renewal, while high *β*-catenin activity triggers differentiation [19, 48]. Accordingly, uniform activation of WNT signalling in the cap mesenchyme (CM) by Sine Oculis Homeobox Homolog 2 (*Six2*)-driven Cre recombinase-mediated expression of stabilised *β*-catenin, or by co-culture with glycogen synthase kinase (GSK) inhibitors results in the uniform emergence of ectopic PTA [27, 32, 45]. Ectopic expression of *Wnt9b* or *Wnt1* (which can substitute for WNT9b) in Six2+ NPCs, on the other hand, results in ectopic differentiation in some, but not all transgenic lines [27, 48]. This likely reflects a concentration-dependent effect as the study that observes differentiation in response to uniform *Six2-Cre*-driven expression of *Wnt9b* reports both renewal and differentiation in response to mosaic *Wnt9b* expression in SIX2+ progenitors [48]. Similarly, cultures of metanephric mesenchyme with WNT9b or WNT1, added either as recombinant protein or secreted from engineered cell lines, yield mixed results, with ectopic differentiation observed in some, but not in all experimental conditions [3, 4].

WNT9b and *β*-catenin counteract the transcription factor SIX2, which supports its own expression, and prevents the differentiation of NPCs by repressing PTA markers such as *Wnt4* and Fibroblast growth factor (*Fgf*)8 [4, 46]. Bone morphogenetic protein (BMP)7 and various FGFs support *Six2* and *Cited1* expression and thereby block PTA formation [3, 15, 61]. In the absence of *Six2*, ectopic renal vesicles form on the dorsal (cortical) side of the ureteric bud (UB), and the NPC pool depletes rapidly, terminating nephrogenesis after induction of only a few nephrons [54]. WNT4 is necessary for PTA formation [59], and by itself sufficient to trigger a mesenchymal-to-epithelial transition and subsequent tubulogenesis, although other members of the WNT family (WNT1, WNT3a, (B) WNT7a, WNT7b, but not WNT11) can substitute for WNT4 [28]. WNT4 and the fibroblast growth factor FGF8 engage in a positive feedback by supporting each other’s expression, but FGF8 supports PTA formation also independently of WNT4, in part by increasing cell motility [47, 55]. Cell tracking revealed that NPCs move largely stochastically in the CM, and cells that initiate *Wnt4* expression can still return to the *Six2* -positive progenitor pool [7, 33, 35]. There is thus no obvious barrier within the metanephric mesenchyme that would limit cells to PTA formation.

How is nephrogenesis then restricted to the branch corner region? Given the complexity of canonical WNT signaling with its many redundant pathway components [29, 39, 49] and the extensive cross-talk of the WNT9b/WNT4/SIX2 core network with other signalling pathways [3, 12], pre-existing differences in WNT responsiveness could, in principle, spatially restrict PTA/RV formation. Also, BMP7 signalling via the mitogen-activated protein kinase (MAPK) pathway maintains progenitors, while SMAD-mediated BMP7 signalling primes progenitors for WNT/*β*-catenin mediated differentiation [2, 3].

Given the stochastic cell movement and the absence of NPC pre-determination [7, 33, 35], any such spatial differences in responsiveness would, however, need to be imposed continuously by external gradients. Given its broad expression in cap mesenchyme and UB [14, 37], it is not obvious how BMP7 could spatially position PTA/RVs to the branch corners. Hitherto unknown secreted factors in the stroma may, however, limit PTA/RV formation to the medullary side [48].

Alternatively, geometric effects can concentrate uniformly secreted proteins in corner regions [43] such that up-concentration of WNT9b could restrict PTA/RV formation to the UB branch corners. Consistent with such a mechanism, mice that lack Mitogen-activated protein kinase kinase (Mek)1 and Mek2 and that have fewer, but elongated UB branches, continue to restrict nephrogenesis to the branch corners and thus display reduced nephrogenesis, even though there is no noticeable *Wnt9b* down-regulation [21, 31]. However, an experimental study has rejected the idea of geometric effects because metanephric mesenchyme abutting an artificially created WNT1 source for two days was found to express a differentiation target gene close to the source and a renewal target gene further away, whether the source was of convex or concave shape [48].

We now revisit the possibility of geometric effects. We combine computational modelling and experiments to show that geometric effects result in higher WNT9b concentration levels in the corner niches between parent and daughter branches hours before PTA/RV form. We further show that NPCs respond to WNT9b in a concentration-dependent manner, and that a local increase in WNT9b induces ectopic PTA/RVs. Using mathematical modelling, we demonstrate that the previous rejection of geometric effects [48] is not supported by the published data, as the expected up-concentration depends not only on the curvature, but also on the gradient length. In fact, we show that the geometry of ureteric branches is optimised to enable the up-concentration of WNT9b in the branch corners. Finally, we show experimentally that PTA/RVs still form in the branch corners when the mesenchyme/stroma is scrambled, making a role of pre-existing patterns in the metanephric mesenchyme and stroma unlikely. We conclude that WNT9b up-concentration via geometric effects provides a simple, robust mechanism to coordinate branching morphogenesis and nephrogenesis during kidney development.

## Results & Discussion

### Geometric effects position nephrons in developing kidneys

We sought to test to what extent geometric effects could concentrate uniformly secreted WNT9b in the corner niches between parent and daughter branches. Using light-sheet microscopy, we obtained the 3D geometry of a developing kidney at embryonic day (E) 12.5, and segmented the UB (green), the outer border of the stroma, the SIX2-positive CM (orange), and the outlines of the PTA/RVs (yellow) (Fig. 1B, Supp. Video S1). The PTA/RVs, as marked by LHX1 (yellow) [42], form in the corners between parent and daughter branches (green). We then simulated the WNT9b concentration profile on the extracted kidney geometry, using the finite element method. We assumed uniform isotropic Fickean diffusion and linear decay, both on the epithelium and in the mesenchyme (for details see Methods section). Based on a comparison of time scales, we model WNT9b as a steady-state gradient. The characteristic time to steady state of a gradient with gradient length λ and decay rate *k* is given by *τ* = (1 + *x*/λ)/2*k* at the readout position *x* [1]. The half-life of WNT9b in the embryonic kidney has not yet been determined, but the turn-over rate of Wingless in the *Drosophila* wing disc has been estimated as 0.0014 s^−1^ [25], in which case *τ* = 12 min at *x* = λ, which is much shorter than the time over which PTA/RVs develop.

For simplicity, we first analysed uniform WNT9b secretion from the epithelium (Neumann boundary condition), even though WNT9b secretion may not be perfectly uniform [4, 26]. We will discuss the impact of non-uniform WNT9b secretion in the next section. The kidney cortex, or the mesenchymal boundaries, are assumed not to constitute diffusion barriers [34]. To visualize the simulated WNT9b concentration gradient in the mesenchyme, we integrated the simulated concentration along rays in normal direction from the UB surface and projected the result back to their origin, where we use a colourmap to display the relative WNT9b concentration (Fig. 1C). Thus, those parts of the ureteric bud that are marked in red are predicted to be adjacent to the mesenchyme where the WNT9b concentration is highest. We find that all PTA/RVs are positioned close to red regions of the ureteric epithelium (Fig. 1C); see the 3D visualisations for closer inspection (Supp. Files S1, S2, Supp. Video S2). The simulated reaction-diffusion model (Methods) only has a single free parameter, the gradient length, λ, which needs to be in the range 20–60 µm to achieve substantial ligand upconcentration in the branch corners (Fig. 1D). Such a gradient length is well within the reported physiological range (λ ∈ [5, 90] µm) [6, 16, 25, 53, 62, 65–67]. For much longer or shorter gradients, the ligand concentrations are either nearly uniformly high or low, respectively.

The average WNT9b concentration in the PTA/RVs is 2-fold higher than in the SIX2 population (Fig. 1E), and in all segmented PTA/RVs, the predicted ligand concentration is higher than the background level in the SIX2 population (Supp. Table 1). We note that the segmented LHX1 domains represent only a rough proxy for the part of the metanephric mesenchyme, where WNT9b induces PTA/RVs, as they include also mature PTA/RVs that have expanded spatially since their first induction by WNT9b [42]. Consistent with this, we find the highest predicted WNT9b concentration in the smallest (i.e., most recently formed) PTA/RVs (Fig. 1F). We further note that the model predicts an elevated WNT9b concentration also at internal branch points, where PTA/RVs are not observed (Fig. 1G). The absence of PTA/RVs in these regions can be accounted to a lack of SIX2-positive CM cells in the inner parts of the developing kidney (Fig. 1G, orange; Supp. Video S2). PTA/RVs therefore cannot form at these internal branch points, even though the local WNT9b concentration is increased.

### The geometry effect is robust to non-uniform *Wnt9b* expression

While *Wnt9b* expression appears largely uniform in the ureteric epithelium, it is excluded from the distal tips, where *Wnt11* is expressed, and its expression level may vary along the ureteric bud [4, 26]. To investigate the impact of such non-uniformity in *Wnt9b* expression, we compared the relative WNT9b upconcentration in PTA/RVs that is obtained with uniform *Wnt9b* expression (Supp. Fig. S1A) with that of four alternative expression patterns: 1) exclusion of *Wnt9b* expression from the distal tips (Supp. Fig. S1B), 2) random, spotty *Wnt9b* expression along the entire ureteric bud (Supp. Fig. S1C), 3) restriction of *Wnt9b* expression to the cortical side (Supp. Fig. S1D), and 4) restriction to the stalk (Supp. Fig. S1E). We obtain a similar level of up-concentration as long as *Wnt9b* expression is not explicitly excluded from the corner region, as is the case when expression is restricted to the cortical side (Supp. Fig. S1F). The effects of spotty expression are smoothed out by diffusion. In conclusion, we find that PTA/RV positioning via geometric effects is rather robust to spatial variations in *Wnt9b* expression.

### Dynamic coordination of branching and PTA/RV positioning

Since PTA/RVs emerge dynamically as the epithelial tree forms via branching morphogenesis, we sought to test the mechanism in time-lapse videos that permit us to follow the coordination between UB branching morphogenesis and the positioning of PTA/RVs over time. Because the imaging of 3D organ cultures is still challenging [10, 18], we took advantage of a liquid-air interface culture system, in which the kidney adopts a flattened shape, allowing the branching process to be followed via live imaging of a 2D projection of the whole tissue. To this end, we cultured E12.25 kidney explants for 48 hours (Fig. 2A). As previously done for the 3D geometries, we segmented the outlines of the UB epithelium, cortex, and PTA/RVs, and solved the same reaction-diffusion equation for WNT9b on the growing 2D geometries (Fig. 2B, Supp. Video S3). Again, the location of the PTA/RVs coincide with the highest ligand concentration, even though the flattened geometry of the cultured explants differs from that in the embryo. Different from before, we do not have a molecular marker of PTA/RV formation in our kidney cultures, limiting us to more mature forms that are discernible by eye. To ensure that the high concentrations are not only correlative, but precede morphological detection of PTA/RVs, we traced the concentration at one location inside PTA/RVs back in time for each individual PTA/RV. (Fig. 2C). Consistent with WNT9b being the most upstream known regulator of PTA/RV formation [4], we predict elevated WNT9b concentrations hours before PTA/RVs can be detected (Fig. 2D). In some cases, our simulations predict high WNT9b concentrations at places where no PTA/RVs form. When we checked these in the 3D renderings of the imaged culture endpoints, we found that these can be accounted to 2D projection artefacts (Fig. 2E, Supp. Fig. S2, Supp. File S2). Even though the cultured kidneys are rather flat, some branches still grow above or below each other, resulting in the false impression of a branch point in the 2D projections.

**Figure 2:**
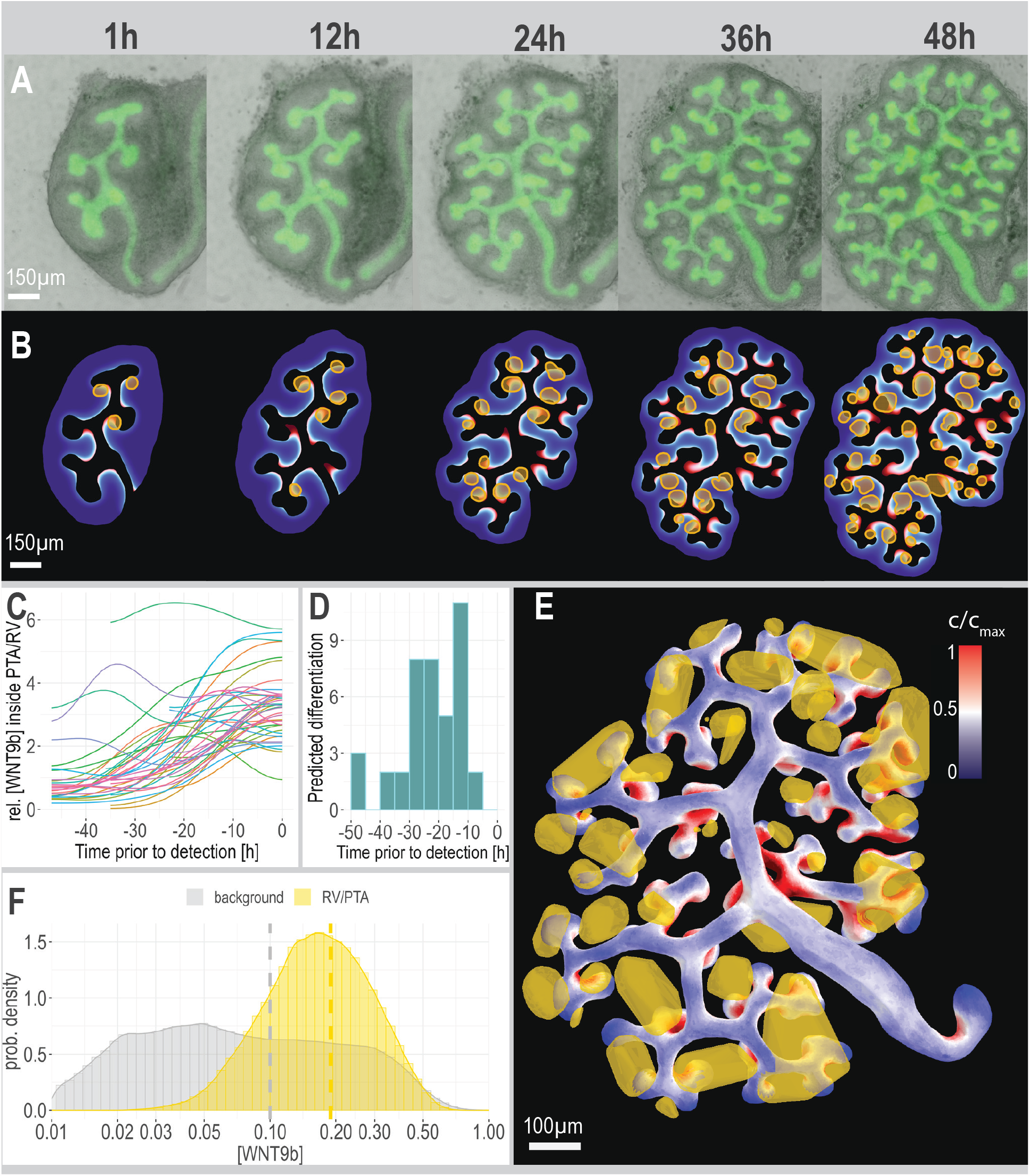
Dynamic coordination of branching and PTA/RV positioning. (A) Widefield live imaging of E12.25 HoxB7-Venus kidneys cultured for 48h; transgenic labeling of the uteric bud (UB). (B) Dynamic simulations of uniform WNT9b secretion from the branching UB yields highest predicted ligand concentration (red) in the UB corner regions, preceding the emergence of PTA/RV (yellow). (C) Simulated WNT9b concentration within PTA/RVs traced back in time, starting from the time point when a PTA-like morphology was first detected in the culture. The concentration profiles were then normalised with control profiles (unit mean concentration of control profiles) and smoothed using a 6-hour Gaussian filter. (D) The predicted concentration profile of WNT9b increases several hours before the first detection of PTA-like morphologies. The time point at which the simulated concentration of WNT9b was first increased by two-fold compared to the control concentration was defined as the time point of NPC differentiation. (E) Simulation of WNT9b on the 3D geometry of the culture endpoint (48h). Uniform secretion of WNT9b from the UB yields the highest predicted ligand concentration (red) in the UB corner regions, coinciding with the positions of PTA/RV (yellow). The plot shows the integrated concentration profile along the normal direction towards the mesenchyme projected back on the surface of the UB. (F) Concentration distribution inside and outside of PTA/RVs from simulations run on the geometry depicted in E. The simulated mean WNT9b concentration inside the PTA/RVs (yellow dashed line; 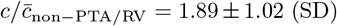) lies significantly above the predicted mean concentration in the mesenchyme (grey dashed line; 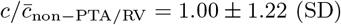); (*p* < 0.001, Welch’s two-sample t-test, *t* = 85.34). All simulations were carried out with λ = 30 µm.

The solution of the steady-state diffusion equation on the 3D tissue geometry confirms that PTA/RVs are located where the predicted ligand concentration is highest (Fig. 2E, Supp. Table S2). As in the 3D embryonic kidney, the WNT9b concentration is predicted to be on average about 2-fold higher in the PTA/RVs (Fig. 2F). Considering that we can only analyse the culture endpoint, we expect to observe nascent nephrons at various developmental stages. Accordingly, large regions exhibiting LHX1 staining are likely indicative of S-shaped bodies. In the cultures, it is well visible how the round PTA/RVs expand to longer structures (Fig. 2B). We conclude that PTA/RV emerge dynamically during branching morphogenesis in the branch corners where the simulations predict high WNT9b concentrations.

### Concentration-dependent response of NPCs to WNT9b

According to our model, the increased WNT9b concentrations in the branch corners induce NPC differentiation into PTA/RVs (Fig. 3A). Consistent with previous reports [32], we confirm a concentration-dependent effect of CHIR99021, a potent GSK inhibitor, on NPC differentiation, as judged by immunostaining for LHX1 and SIX2 to mark PTA/RVs and the CM, respectively (Fig. 3B; shown are segmentation masks of the stainings). While developing nephrons are located in UB corners in controls, uniformly supplied CHIR99021 induces ectopic NPC differentiation. With a lower CHIR concentration of 5 µM, differentiated cells are found below the cortex after 40h, regularly interspaced with SIX2-positive progenitors, whereas no LHX1 is detected close to the UB. A high CHIR concentration of 10 µM rapidly (within 24 hours) induces LHX1 in a large portion of the nephrogenic mesenchyme, which appears to be expanded relative to the UB. In both cases, branching morphogenesis is impaired as GDNF-secreting cap mesenchyme progenitors differentiate. On the other hand, we observe no ectopic differentiation when culturing kidney explants with recombinant WNT9b, even at concentrations as high as 2 µg/ml (data not shown). The observed difference between WNT9b and CHIR could either reflect a low activity of the recombinant WNT9b protein, barriers to its spreading, or the requirement for other signalling factors in addition to WNT9b that are present only in the branch corner and that CHIR bypasses.

**Figure 3:**
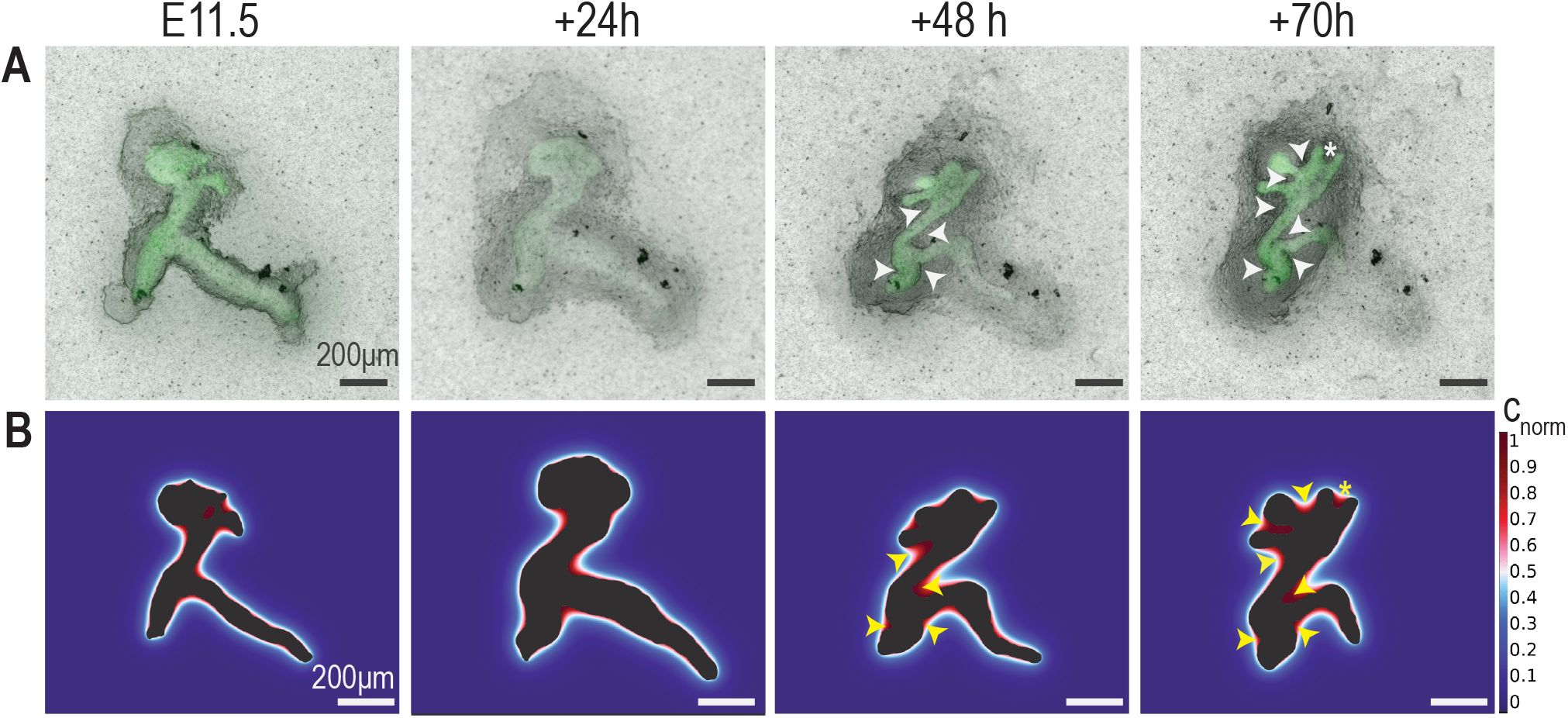
Concentration-dependent response of NPCs to WNT9b. (A) Diffusion simulation on a simplified UB branch, illustrating ligand upconcentration and PTA/RV positioning by geometric effects. (B) 3D Imaris surfaces of HoxB7/myr-Venus kidney explants (segmented UB in green, outer surface transparent), immunostained for LHX1 and SIX2 (control and 5 µM). Control; PTA/RVs form in the corners of UB branches; treated with 5 µM CHIR99021 for 40h; LHX1+ clusters form away from the UB, interspersed with SIX2+ progenitors; treated with 10 µM CHIR99021 for 24h; High uniform WNT activation results in *Lhx1* expression in most of the expanded metanephric mesenchyme. Scale bars 100 µm. (C–E) Concentration-dependent response of Six2-positive primary NPCs, isolated from E12.5 kidneys, to recombinant WNT9b, added either in the medium (C) or on beads (E), as judged by qPCR of differentiation markers (*Lef1, Wnt4*) and renewal markers (*Cited1, Pax8*) (D). At high WNT9b concentrations, expression of *Cited1, Pax8* was no longer detectable (n.d.). (F–L) HoxB7/myr-Venus kidney explants were cultured with control (G) or WNT9b-soaked (H, I) beads (dashed white circles). White arrowheads mark the position of renal vesicles that are forming in close proximity to the WNT9b source. (I) UB branches (black lines) bend toward the WNT9b source over time, potentially following CM cells that are attracted to the WNT9b-soaked bead. Ectopic branching from the future ureter (white asterisk). (J–L) 3D rendered LSFM data of the explant culture endpoints, immunostained for LHX1 and SIX2 for control (G) and WNT9b-soaked (H, I) beads. The white asterisk marks the same branch as in I.

To test whether we used a sufficient concentration of recombinant WNT9b, we isolated SIX2+ NPCs from E12.5 kidneys using fluorescent-activated cell sorting (FACS) and exposed them for 18 hours to different concentrations of recombinant WNT9b (Fig. 3C). We subsequently used qPCR to measure the fold change in the expression of differentiation (*Lef1, Wnt4*) and renewal (*Cited1, Pax8*) markers (Fig. 3D). An increase in *Wnt4* and *Lef1* expression is observed at 0.6 and 1 µg/ml recombinant WNT9b respectively, and plateaus for higher WNT9b concentrations. The renewal markers *Cited1* and *Pax8* where expressed at low level and fell below the detection level (n.d.) at high WNT9b concentrations. This shows that NPCs are, in principle, responsive to the concentrations of recombinant WNT9b used in the explant cultures.

To test whether the lack of a response of NPCs to soluble recombinant WNT9b in explant cultures was due to limitations in its spreading, we next used WNT9b-soaked beads. The beads allow us to deliver WNT9b to the top of the kidney explant where it is rapidly engulfed, while the culture medium is located below the filter at the bottom of the culture (Fig. 3F). The beads were soaked in 40 µg/ml recombinant WNT9b and elicit similar level of *Wnt4* expression in the isolated NPCs as 0.6–2 µg/ml soluble WNT9b, while *Lef1* expression is considerably higher with the beads (Fig. 3D). We next co-cultured E11.5 kidney explants with either control or WNT9b-soaked beads (Fig. 3G–I, Supp. Movie 5), and confirmed PTA/RV and nephron progenitor positions by immunostaining culture endpoints for LHX1 and SIX2 (Fig. 3J–L). PTA/RV formation could indeed be observed close to the WNT9b source (Fig. 3K–L, Supp. Fig. S3). Taken together, these results show that high WNT9b concentrations trigger differentiation also outside the branch corners.

We note that in one example, the most proximal branches and the future ureter bent towards the WNT9b source during explant culture (Fig. 3L, Supp. Movie 5), possibly as a consequence of the attraction of GDNF-secreting CM towards the bead. GDNF-secreting CM close to the bead could also cause ectopic branching from the future ureter, which is associated with a developing nephron (Fig. 3H,I, Supp. Movie 5). Such ectopic branches are sometimes also observed in regular kidney cultures [63].

During extended culture periods, PTA/RVs begin to emerge at the top of branching ureter tips (Supp. Fig. S3A,D–F). 3D simulations, based on the culture endpoint geometries (Supp. Fig. S3B), demonstrate that the PTA/RVs form precisely at the positions where high WNT9b concentrations are predicted to occur as a result of a zero-flux boundary condition that arises at the tissue-air interface because WNT9b cannot diffuse into the air (Supp. Fig. S3D–F). Notably, when kidneys are cultured in different conditions, where they are surrounded by liquid, they tend to develop more branch points and do not form or delay the formation of these culture-induced PTA/RVs [52].

In contrast, the ectopic PTA/RVs that are induced by WNT9b-loaded beads form at a distance from the UB, where in the absence of the bead, low WNT9b concentrations are predicted (Supp. Fig. S3B,C,G,H). We note that the culture-induced PTA/RVs are smaller than the bead-induced PTA/RVs. As PTA/RVs increase in volume over time, as also visible in our live imaging experiments (Supp. Fig. S2, Supp. Movie S5), the PTA/RV volume can serve as a proxy for developmental age (Supp. Fig. S3I). The smaller culture-induced PTA/RVs thus tend to develop later in the culture process, typically when the tissue flattens. In contrast, the larger bead-induced PTAs form around the same time as the first endogenous PTAs in the UB corners.

In summary, we reproduce earlier reports that soluble WNT9b fails to induce ectopic PTA/RVs in explant cultures. Similar concentrations of recombinant WNT9b, however, induce *Wnt4* expression in isolated NPCs, suggesting that no additional pre-pattern is required to enable NPCs to respond to WNT9b. The fact that bead-loaded WNT9b induced differentiation both in isolated NPC cultures and in explant cultures rather suggests that the soluble WNT9b provided in the medium fails to reach SIX+ NPCs by diffusion to a sufficient level. We further show that at late culture time points, ectopic PTA/RVs emerge exactly at those positions where the model predicts high WNT9b concentrations due to a zero-flux boundary condition at the tissue-air interface. Together, this supports our proposal that a local increase in the WNT9b concentration due to geometric effects restricts nephrogenesis to the branch corners (Fig. 3A).

### PTA/RVs continue to form in the branch corners when mesenchyme and stroma are scrambled

PTA/RVs have been proposed to be positioned by pre-existing patterns that lead to increased WNT responsiveness in the branch corners [48]. To date, no such pre-pattern has been identified. We sought to test this possibility by culturing ureteric buds with scrambled kidney mesenchyme (Fig. 4A). While we cannot exclude rapid cell sorting based on differential adhesion, the continued restriction of PTA/RVs to the branch corners would lend further support to our proposal of the role of geometric effects, while making a role of pre-patterning less likely. Live imaging of the recombined cultures over 70 hours reveals the continued restriction of PTA/RV emergence to the branch corners (Fig. 4A, white arrows), as judged by their morphology. As always, the emergence of morphologically visible PTA/RVs is delayed relative to the predicted emergence of high WNT9b signalling, as can be seen by following the emergence of PTA-like morphologies relative to the predicted levels of WNT9b (Fig. 4B). As such, we expect that a PTAs would still have formed in the one position where our model predicts high WNT9b concentrations, but no PTA-like morphology can be observed by 70 hours (Fig. 4B, asterisk). In summary, the continued restriction of PTA/RVa to branch corners, despite the scrambling of the mesenchyme, suggests that a mesenchymal/stromal pre-pattern is not necessary for the positioning of PTA/RVs.

**Figure 4:**
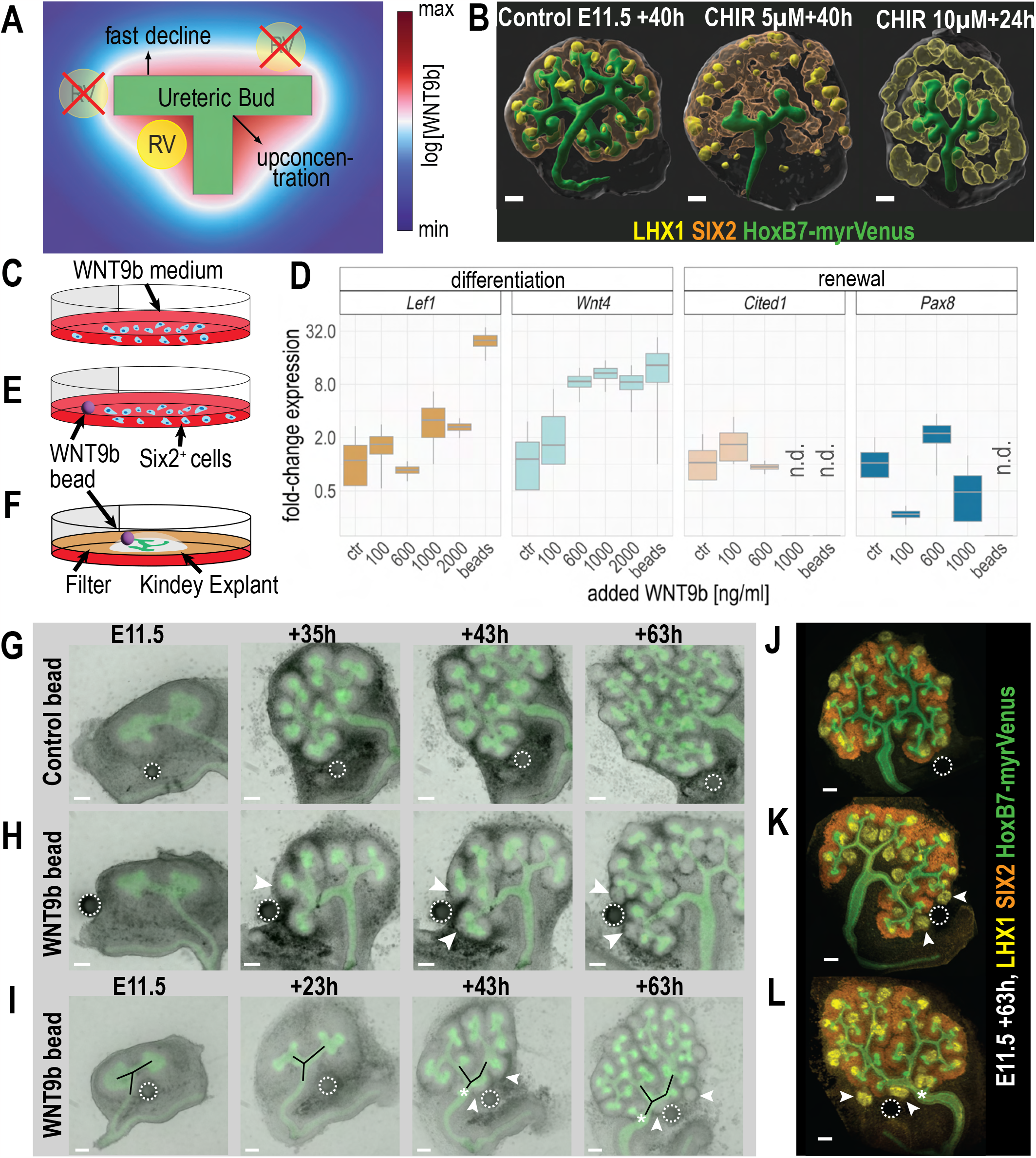
PTA/RVs continue to form in the branch corners when mesenchyme and stroma are scrambled. (A) Kidney explant cultured with scrambled mesenchyme. The metanephric mesenchyme of multiple E11.5 kidneys was separated from the ureteric epithelium, lysed, and added back to a cultured epithelial bud. After 48 h, PTA-like morphologies became visible in the corner regions (white arrows). (B) The UB was segmented and used to predict the WN9b concentrations with a steady-state diffusion model. In the simulations, the UB boundary serves as a uniform source of WNT9b via Neumann boundary conditions. Regions with high relative concentration (red) co-localize with locations where we observe PTA-like morphologies (yellow arrows), except in one position (asterisk), where the emergence of PTA-like morphologies is likely delayed.

### Optimal branch kink angle and curvature

We wondered whether the kidney geometry was particularly suited to permit the coordination of branching morphogenesis and nephrogenesis via geometric effects. The ligand upconcentration can be expected to depend on two geometric properties: the branching angle, and the tissue curvature in the corner.

To investigate the impact of the branching angle, we simulated the steady-state reaction-diffusion equation (Methods) on a 2D domain with a 1D ligand source of constant length, *L*, kinked in the middle by an angle Θ (inset Fig. 5A). As expected, we find the ratio of the resulting concentration in the corner, *c*_1_, to the concentration at the outer ends of the source, *c*_2_, to monotonically increase with smaller Θ (Fig. 5A). The results are similar whether we use constant uniform production or a constant outflux from the source, and are independent of the relative gradient lengths λ/*L*. For the smallest tested angle, Θ = 10°, we find a more than 30-fold higher concentration at the kink. However, for such a sharp kink, branches would grow into each other upon consecutive branching, resulting in rapid termination of branching. To permit continued branching, the branching angle must exceed 90° on average. In the E12.5 kidney, we observe the angles between tip and parent branches to range from 81–121°, with a mean of 98.8 *±* 11.9° (SD) (Fig. 5B). This value is slightly lower than previously reported, possibly because previous analyses included internal angles, which increase during development as the ureteric tree remodels [56–58]. For the measured branching angles (Fig. 5B), we expect a 2-fold upconcentration of the ligand in the corner niches (Fig. 5A), as indeed observed in our E12.5 kidney simulations (Figs. 1E, 2F). We conclude that the branching angles in the kidney maximise the WNT9b concentration in the branch corners, while still permitting regular branching of most tips.

**Figure 5:**
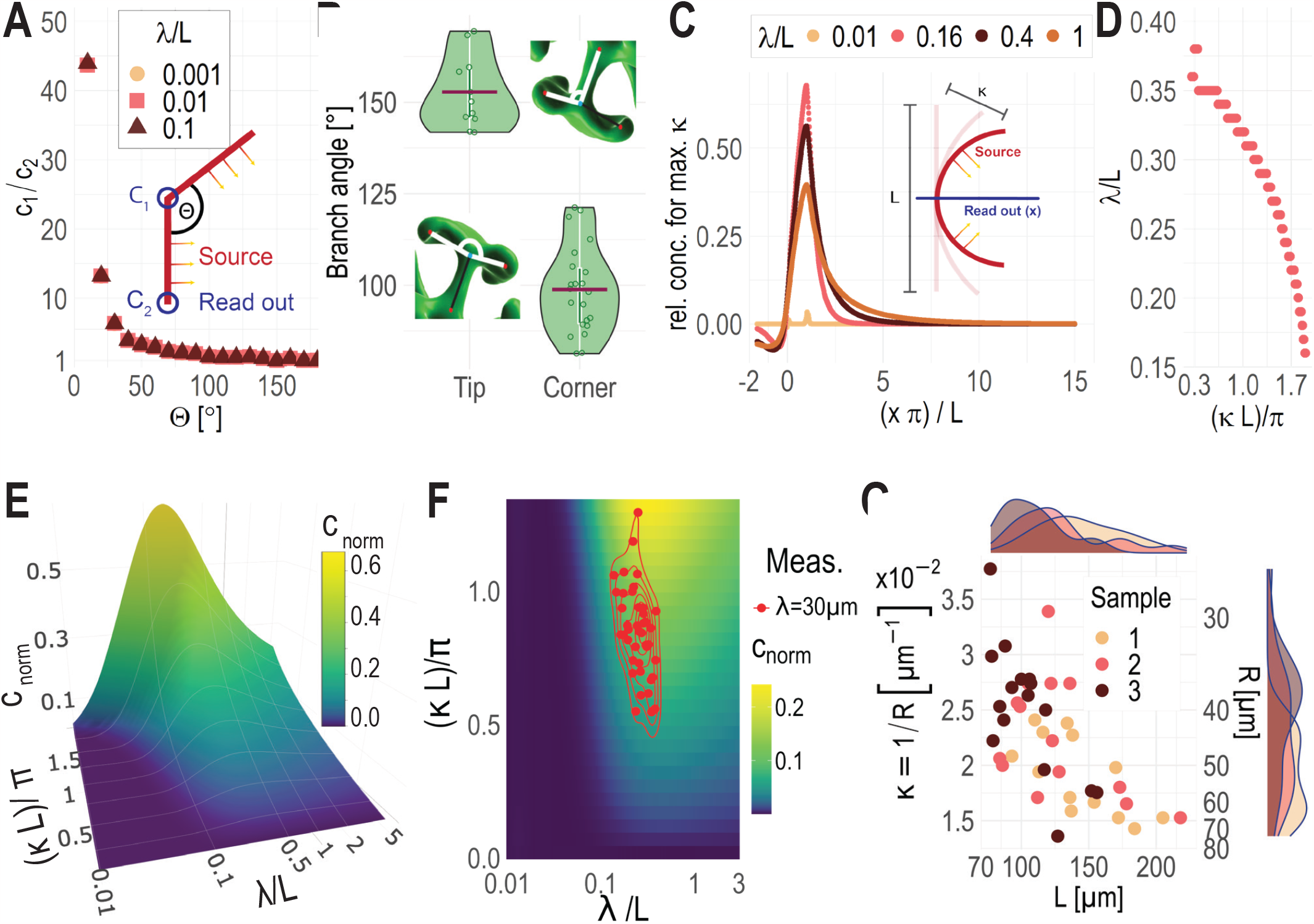
Optimal source kink angle and source curvature. (A) The concentration at the corner, *c*_1_ is highest relative to the endpoint of the source, *c*_2_, the smaller the angle, Θ, between the source parts. The same results are obtained with different relative gradient lengths, λ/*L*. The 2D simulations were carried out with constant flux boundary conditions at the source, but similar results are obtained with constant production in the source (inset, red). Note the logarithmic axes. (B) Measured branching angles in embryonic kidneys (E12.5). Measured were the outer angle (tip) and inner angle (corner). The insets describe the location of measurement. (C) The concentration profile originating from the center (inset, blue) of half-circular sources (inset, red) depend on the relative gradient length λ/*L*. The highest concentrations are observed in the concave part for λ/*L* = 0.16. (D) The highest ligand upconcentration depends on relative gradient length, λ/*L* and the curvature of the source, *κ*. The higher the curvature, the lower, the optimal gradient length for which maximal upconcentration is achieved. Displayed are the values with the highest concentration per simulated curvature, i.e., the ridge in panel E. (E) Relationship of relative gradient length, λ/*L*, curvature of the source, *κ*, and the relative concentration increase, *c*_norm_ at the readout position *x* = *L*/*π*. The concentration increases with curvature and peaks at λ/*L* = 0.16. (F) For λ = 30 µm, the measured branch lengths and curvatures in E12.5 kidneys (panel G) result in maximal upconcentration (red dots). (G) Measurements of curvature, radius, and length of ureter branch corners at positions where renal vesicles form. Analysis is based on three E12.5 kidneys.

To investigate the impact of the branch curvature, we consider a bent 1D source (red line) of length *L* and curvature *κ* = 1/*R*, where *R* is the radius of the circular arc that the source follows (inset Fig. 5C). As the investigated geometric effects must be independent of the chosen length scale, we normalise all length scales with the source length, *L*. Accordingly, we consider a normalised gradient length, λ/*L*, a normalised curvature, *κL*/*π*, and a normalised distance from the source, *xπ*/*L*. Here, *L*/*π* is the radius of a semi-circle with circumference *L*. Moreover, we consider a normalised concentration that quantifies the relative difference between the concentration profile resulting from a curved (*c*(*κ, x*)) and straight (*c*(0, *x*)) source, normalised by the concentration at the straight source, *c*(0, 0):

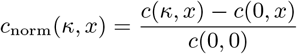

We analyse the concentration profile along a straight line (blue) orthogonal to the center of the source (Fig. 5C, inset). Concave curvature (*x* > 0) results in ligand upconcentration, while convex curvature (*x* < 0) has the opposite effect and results in reduced concentrations (Fig. 5C). The normalised relative concentration profiles for maximal curvature (*κ* = *π*/*L*) peak at a similar relative distance from the source, *x* ≈ *L*/*π*, with longer gradients (larger λ/*L*) peaking slightly closer to the source (Fig. 5C). The highest upconcentration is achieved for λ/*L* = 0.16. At smaller curvatures, the optimal relative gradient length increases (Fig. 5D), and the maximal upconcentration drops (Fig. 5E). Fig. 5D traces the ridge of the surface shown in Fig. 5E.

While the strongest upconcentration is achieved for the greatest curvature (Fig. 5E,F), high curvature implies small corner radii that may not leave sufficient room to accommodate PTA/RVs. The smallest PTA/RVs that we detect in E12.5 kidneys have a diameter of about 25 µm, such that the curvature cannot exceed 0.08 µm^−1^. The larger PTA/RVs in E12.5 kidneys reach 100 µm in diameter along the UB, which bounds the curvature to about 0.02 µm^−1^ from below, if spherical PTA/RVs are to fill the niche without any gaps. We note that at later stages, PTA/RVs, however, depart from a spherical shape and develop into S-shaped bodies.

To determine the physiological branch length and corner curvature, we obtained three E12.5 kidneys and measured the curvature and length of each UB where a PTA/RV was detected (Fig. 5G). The curvature of the branch corners is between 0.014– 0.038 µm^−1^ (Fig. 5G). We observe an anti-correlation between the branch length and branch curvature, suggesting that the curvature is highest in newly forming PTAs. This is consistent with previous observations that showed that the expanding PTA/RV reduces the curvature of the ureteric epithelium [35].

For a given curvature, *κ*, and source length, *L*, there is an optimal gradient length, λ, that yields the highest upconcentration in the branch corner (Fig. 5E,F). For the measured branch curvatures (*κ* = 0.0224±0.0056 µm^−1^ (SD)) and lengths (*L* = 123.65±35.08 µm), we predict the highest ligand upconcentration for a gradient length of λ = 30 µm (Fig. 5F). This is consistent with our simulations on the 3D embryonic kidney geometries (Fig 1D), and such a gradient length is well within the reported range for morphogen gradients in other developmental systems, i.e. λ ∈ [5, 90] µm [6, 16, 25, 62, 65–67].

We conclude that the geometry of the ureteric tree appears to be optimised to permit a substantial upconcentration of WNT9b at the branch corners over large enough distances to induce sizeable PTA/RVs, while permitting continued branching during development.

### Previous rejection of geometric effects is unwarranted

Finally, we revisit a previous study that had rejected the possibility that the WNT9b concentration is elevated in the branch corners by geometric effects [48]. In the study, *Wnt1* was expressed in stromal cells of isolated metanephric mesenchyme and the target gene expression was analysed relative to the morphology of the WNT source (Fig. 6A). The authors ruled out geometric effects because they observed the expression of the differentiation marker, *C1qdc2*, directly abutting to the source, irrespective of whether the source had a concave or convex shape. However, when we simulate uniform WNT1 secretion from the reported source shape, we predict very similar response patterns as reported (Fig. 6B). The lack of visible differences between the convex and concave parts (Fig. 6C) can be accounted to the relatively low curvature of the artificial source (Fig. 6D), which leads to a comparably low up-concentration for the likely WNT9b gradient length of about 30 µm (Fig. 6E). To observe noticeable geometric effects, the gradient length would need to be considerably larger, but also then, the strongest difference between convex and concave boundaries would only be found at a considerable distance from the source (Fig. 6F). The uniform response that is observed along the artificial WNT1 source likely reflects over-expression of *Wnt1* compared to endogenous *Wnt9b* expression in the ureteric bud. The misinterpretation of the reported experiment demonstrates the importance of computational modelling in the analysis of patterning mechanisms.

**Figure 6:**
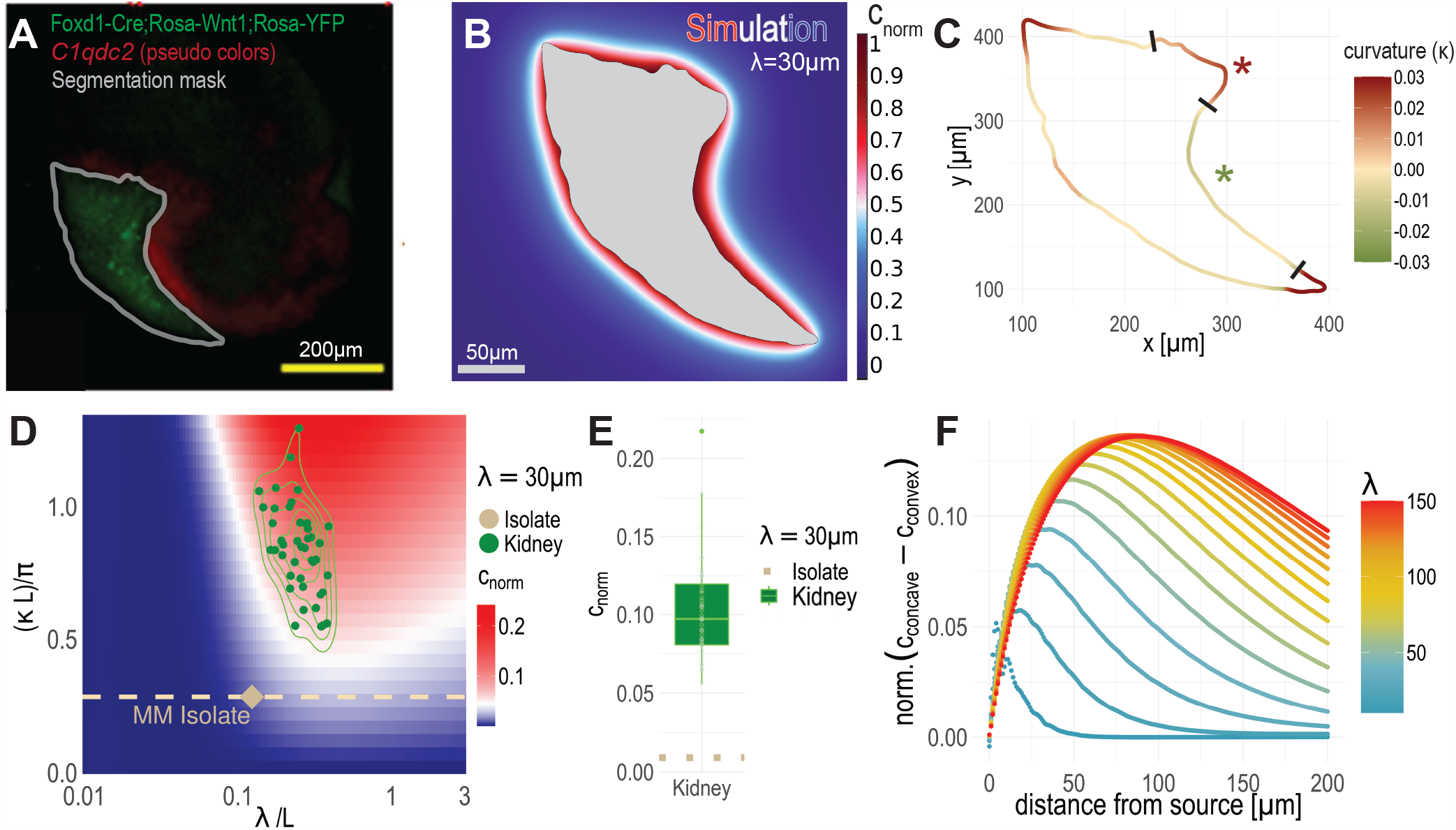
Reassessment of a previous experiment evaluating the impact of geometric effects on WNT signalling in the kidney. (A) Ramalingam *et al*. cultured E11.5 Foxd1-Cre;Rosa-Wnt1;Rosa-YFP isolated metanephric mesenchyme for two days and analysed the expression of a differentiation target gene (*C1qdc2*) (red) with in situ hybridisation relative to the WNT1-secreting source (green). The panel was reproduced from [48]. The boundary (grey) of the WNT1-secreting source (green) was added for illustration. (B) The predicted relative steady-state concentration when WNT1 is secreted uniformly (Neumann boundary conditions) from the segmented boundary of the source (panel A) for a gradient length of 30 µm. (C) The curvature of the WNT1 source boundary in panel A. Concave sections are green, convex sections red. The NPC-facing sections with the highest curvature (marked with asterisks) were extracted for further analysis. (D) The concave part extracted in panel C has a smaller curvature (dashed line) than all branches measured in E12.5 kidneys (green). (E) Given the small curvature, the predicted up-concentration (dots) is negligible compared to that predicted for ureteric buds (green). (F) To achieve meaningful concentration differences between the concave and convex parts of the isolate, unphysiologically large gradient lengths (red) would be required.

## Conclusion

During kidney development, nephrogenesis and branching morphogenesis must be coordinated such that a single nephron connects to each ureteric branch point, thereby ensuring efficient drainage of the kidney. How this coordination is achieved has long remained elusive, even though the key molecular regulators, most notably WNT9b, have long been identified [4]. We have now shown that geometric effects lead to a higher WNT9b concentration in the branch corners of the ureteric tree, even though *Wnt9b* is expressed largely uniformly along the ureteric epithelium. As WNT9b induces PTA/RVs, these geometric effects can coordinate nephrogenesis with branching morphogenesis.

We show that the geometric effects are strongest, the smaller the branch angle, and the higher the branch curvature (Fig. 5A,C). The branch angle is indeed the smallest that it can be while permitting continued branching, and the corner curvature is the highest it can be while accommodating the PTA/RVs in the branch corners. This suggests that the geometry is optimised to permit strong ligand upconcentration in the branch corners. Finally, we show that the ligand upconcentration depends on the gradient length, which must be in a physiologically plausible range to ensure significant ligand upconcentration. While the WNT9b gradient length remains to be determined experimentally, we expect it to be at least 20 µm.

The role of tissue geometry for patterning has long been recognised [41, 43], and it will be interesting to see to what extent the here-identified patterning principle will apply to other developmental processes. The geometric concentration effect could be exploited also in tissue engineering applications [44] to generate non-uniform, localised morphogen distributions. Beyond the geometry, also the tissue boundary can play a role. Thus, the emergence of ectopic PTA/RVs after extended culturing of E11.5 kidney explants can be explained with the up-concentration of WNT9b at the zero-flux boundary at the tissue-air interface of the flattened explant (Supp. Fig. S3D–F). At late stages of kidney development, PTA/RVs are found to emerge also on the cortical side [50]. It will be interesting to unravel what changes lead to this change in location.

### Limitations of the Study

Direct imaging and reliable quantification of morphogen gradients in tissues and organs remains technically challenging, preventing us from directly quantifying WNT9b protein concentrations in the developing kidney. We are thus restricted to make conclusions about the protein distribution through our simulations, assuming constant diffusion and degradation in steady state. Also, we could not measure the effective gradient length *in vivo* and estimated it based on physiological values in the literature and best fits in our simulations. Additionally, 3D imaging and simulations become increasingly difficult with increasing size and complexity of the tissue. We therefore restricted our study to the early stages of kidney development, and we could not study the extent of geometric effects at later stages. As the light-sheet imaging of 3D kidney cultures remains challenging [18], we followed the branching process via live imaging of a 2D projection of flattened embryonic kidney explants in a liquid-air interface culture system. While flat, some important 3D effects are invisible. Moreover, as we did not have a genetic marker of emerging PTA/RV, we were limited to the morphological detection of their formation.

## Supporting information

Supplementary 3D visualisations

## Acknowledgements

This work was funded by the Swiss National Science Foundation under Sinergia grant no. CRSII5 170930. We thank Marco Meer for interesting discussions.

## Author Contributions

Experiments: L.C., M.M., N.D; Image processing: M.M., L.C.; Simulations: M.M.; Theory and modelling: D.I., R.V., M.M.; Writing: D.I., M.M., L.C., R.V., N.D.; Visualisation: M.M., L.C., R.V.; Supervision: R.V., D.I.; Conceptualisation: D.I.; Project administration: D.I.; Funding acquisition: D.I.

## Methods

### Data and Code Availability

The source code and COMSOL model files are released under the 3-clause BSD license and are available as a public git repository at https://git.bsse.ethz.ch/iber/Publications/2022_mederacke_conrad_renal_vesicles.

### Experimental Model

#### Ethical statement

Animal experiments were performed in accordance with Swiss federal law and the ordinance issued by the Canton Basel-Stadt and approved by the veterinary office of the Canton Basel-Stadt, Switzerland (approval number 2777/26711).

#### Mouse strains

Labeling of the ureteric bud was achieved by using the Hoxb7/myr-Venus transgenic allele [MGI: Tg(Hoxb7-Venus*)17Cos; [5]]. Timed pregnancies were set and checked daily for vaginal plugs to obtain the desired embryonic ages. Here, homozygous Hoxb7/myr-Venus males were crossed with RjOrl:SWISS wild-type females (higher pregnancy rates and larger litters).

Labeling of the NPC population was achieved by using the Six2TGC conditional knock-in/knockout mouse line [MGI: B6;129-Six2tm3(EGFP/cre/ERT2)Amc/J [30]]. To combine Hoxb7/myr-Venus and Six2TGC labeling, heterozygous males (Six2TGC/Hoxb7) were mated with homozygous Ai14 [MGI: B6.Cg-Gt(ROSA)26Sortm14(CAG-tdTomato)Hze/J [38]] females. Expression of the fluorophore was induced by intraperitoneal injecting the pregnant females with 200 µl of 10 µg/ml tamoxifen 24h before collection of the embryos.

#### Animal housing and husbandry conditions

All mice included in the study exhibit a healthy phenotype and have not been utilized in any prior experiments. Housing and husbandry were performed after the guidelines and with the protocols of the veterinary office of the Canton Basel-Stadt, Switzerland.

### Method Details

#### Primary cell isolation and experiments

E11.5 kidneys from Six2TGC+ embryos were dissected and transferred in an Eppendorf tube containing DMEM-F12(Gibco) on ice. Once all the kidneys were collected, DMEM-F12 was substituted by 500 µl of trypLE (Gibco). Kidneys were treated for 5’ at 37°C with trypLE, followed by repetitive pipetting. After trypsin treatment, cells were centrifuged for 3’ at 800g. The pellet was resuspended in PBS supplemented with 2% FBS (Gibco) and 10mM EDTA (Thermo). Cells were filtered using a 40 µm nylon cell strainer and were kept on ice until FACS sorting. The Six2+ cells were directly FACS sorted into a 96 well plate supplemented with 10% FBS/ Dulbecco’s modified Eagle’s medium (DMEM,Gibco) in the presence of 4 µM DMSO (Thermo). After 24h the cells were treated with WNT9B (R&D, 3669-WN) as indicated. RNA from each experimental condition was isolated by using Absolutely Rna Nanoperp kit (Agilent technologies). CDNA was generated with PrimeScript kit (Takara). SYBR Green SuperMix (Bio-Rad) was used for qPCR reactions using 3 µL of diluted cDNA (1 mg RNA equivalent in 100 mL). GAPDH was used as the reference gene. QuantStudio 3 (Thermo Fisher Scientific) instrument and software were used to determine relative gene expression levels using the delta-delta Ct method. Primer sequences were manually designed or were obtained from prior publications.

mouse gapdh forward AACTTTGGCATTGTGGAAGG

mouse gapdh reverse ATCCACAGTCTTCTGGGTGG

mouse pax 8 forward CACAAAGGCCCCTCCTAGTT

mouse pax 8 reverse GCGAGTGTCCCTCAGTCTGT

mouse wnt 4 forward CTCAAAGGCCTGATCCAGAG

mouse wnt 4 reverse TCACAGCCACACTTCTCCAG

mouse cited 1 forward CATCCTTCAACCTGCATCCT

mouse cited 1 reverse ACCAGCAGGAGGAGAGACAG

mouse lef1 forward TCATCACCTACAGCGACGAG

mouse lef1 reverse GAAGGTGGGGATTTCAGGAG

#### Explant culture live imaging

Embryonic kidneys were dissected using fine forceps and tungsten needles in cold phosphate-buffered saline (PBS) and collected in a petri dish containing cold culture medium (DMEM-F12 supplemented with 10% fetal bovine serum, 1x GlutaMAX, 1x penicillin/streptomycin). The dissected kidneys were cultured at liquid-air interface on top of a porous filter membrane insert in a 6-well plate containing 1.5ml culture medium per well. To activate WNT signalling, the medium was supplemented with 5 or 10mM CHIR99021 (Stemcell technologies 72052), or with 200, 500, or 750ng/ml WNT9b (Peprotech 120-49) diluted in the culture medium. To locally deliver WNT9b, Affi-Gel Blue beads (Biorad 153-7302) were rinsed with PBS and incubated in 40µg/ml WNT9b (Peprotech 120-49) or in PBS (control) for 1 h at 37°C. Several beads were transferred into PBS to rinse off excess protein solution, one bead was selected using a P10 micro-pipette, and positioned close or on top of a kidney explant using fine forceps. The culture medium was changed every 48h.

#### Volumetric lightsheet microscopy and image analysis

Tissue clearing, immunofluorescence, and lightsheet fluorescence microscopy (LSFM) were performed as previously described [10, 60]. Blocking and antibody incubation was done using 3% bovine serum albumin (BSA) (Sigma-Aldrich A7906-10G) and 0.3% TritonX (Sigma-Aldrich T8787) in 1x PBS. Primary antibodies were incubated for 48h using the following dilutions: anti-LIM1/LHX1 (abcam ab229474) 1:200; anti-LIM1/LHX1 (antibody 4F2, deposited to the DSHB by Jessell, T.M. / Brenner-Morton, S. (DSHB Hybridoma Product 4F2)) 1:50; anti-SIX2 (proteintech 11562-1-AP) 1:200. All secondary antibodies were diluted 1:500 and incubated for 24h. The volumetric LSFM imaging data was imported into Imaris (Bitplane, Oxford Instruments) and 3D-rendered. To obtain segmentations of the ureteric bud, the renal vesicles, and the kidney cortex, Imaris surfaces were generated by intensity- and volume-based thresholding of the respective channels and manually corrected where needed (DAPI nuclear staining was imaged using a 405 nm laser, myr-Venus was imaged using a 488 nm laser, fluorophore-conjugated secondary antibodies were imaged using a 561 nm or a 647 nm laser). For the branching angle measurements, binary images were created in Imaris, skeletonised in 3D using the Fiji plugin “Skeletonize3D” and pruned over several iterations to remove wrongly segmented, small side branches. The cleaned tree was analysed using BoneJ2 to measure global angles at triple-junctions [13].

#### UB cultures with scrambled mesenchyme

At E11.5, kidneys were dissected as described above. The tissues were then treated with trypLE (Gibco) for 8 minutes at 37°C to weaken the epithelial-mesenchymal connection, and subsequently returned to ice-cold DMEM-F12 (Gibco). Using a fine forceps, the two cell layers were carefully separated, after which the mesenchymal cells were subjected to a further 3-minute digestion before being dissociated into a single-cell suspension via repetitive pipetting. The mesenchymal cells were then centrifuged for 3 minutes at 800g to halt the enzymatic treatment, and the resulting pellet was resuspended in a small volume of PBS. Ureteric epithelial structures were placed onto a porous filter, overlaid with the mesenchymal cell suspension, and cultured for 72 hours using the methods described previously.

#### Computational model

We describe the spatio-temporal distribution of WNT9b, 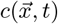, with a steady-state (*∂*c/*∂*t = 0) reaction-diffusion equation of the form

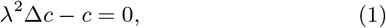

where Δ is the Laplace operator. The equation has a single free parameter, λ, the gradient length, which is determined by the ratio of the diffusion coefficient and the turn-over rate. As the WNT gradient length has not yet been reported in mice, we set the gradient length to our estimated optimal λ = 30 µm (Figs. 1D, 5F), unless otherwise stated. On the surface of the ureter, we used a Neumann boundary condition of the form

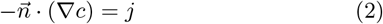

with the surface normal vector 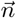. As the absolute WNT9b concentration is not known, we only consider relative concentrations, c/c_max_, and j can be chosen freely without affecting our conclusions. The kidney cortex, or the mesenchyme boundaries are unlikely to constitute diffusion barriers in uncultured kidneys [34], and we therefore solved the equations in a large enough box that the outer boundaries do not affect the solution.

We used the same simulation setup for the 3D and 2D kidney simulations. The 2D simulations differ only in that the reaction-diffusion equation is solved on a growing domain. As the cultured kidneys were imaged only every 60 minutes to avoid photo-toxicity, we interpolated the time points in between by computing displacement fields describing the morphological change from one captured culture time point to the next, using minimal distance mapping, as described earlier [51]. These vector fields are implemented in the simulation to prescribe a mesh deformation mimicking the observed growth process and allowing to predict WNT9b localisation on a continuous growing domain (Supp. Movie. S4) [17, 40].

All simulations were performed using the finite element method in COMSOL Multiphysics v6.0 (COMSOL AB, Stockholm, Sweden), as described before [17, 40].

#### Tracking simulated concentration over time

To monitor the temporal changes in concentration within live image data of kidney cultures, point probes were strategically positioned at locations and time points where PTA/RV were detected. Subsequently, these probes were traced back over time. The resulting intensity profiles were smoothed using a Gaussian kernel spanning a 5-hour window and normalized with a control concentration profile. The control profile was generated by randomly placing five point probes at the boundary of the UB outside of any identified PTA/RV (Fig. 2D). The time point at which the simulated concentration of WNT9b was first two-fold increased compared to the control concentration was predicted to correspond to the initiation of stem cell differentiation.

#### Curvature inference from 2D contours

To calculate the curvature of 2D images (see Supplementary Figure S2C), we fitted circles through each pixel and its two evenly spaced neighbors on both sides, using a distance of 100 µm. If the circle’s center fell inside the segmentation, we assigned a positive sign (indicating convex curvature); otherwise, we assigned a negative sign (indicating concave curvature).

### Additional Resources

2D plots were generated using R and ggplot2 [64], 3D plots were generated using python and plotly [22]. Meshes were visualized and organised using Blender 3.1 [9].

## Supplementary Material

### Supplementary Data

**Table S1:**
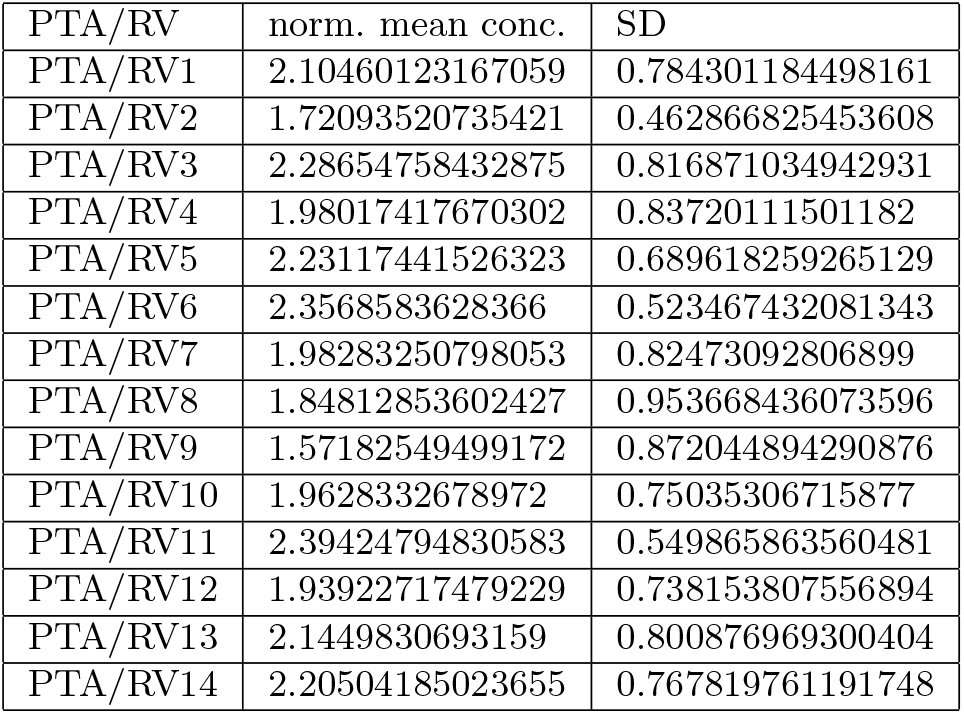
Predicted concentration in the PTA/RVs of an E12.5 kidney. Simulated normalized mean concentration, 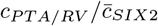, inside each segmented PTA/RV and standard deviation. From the E12.5 Kidney used for Fig.1D.

**Table S2:**
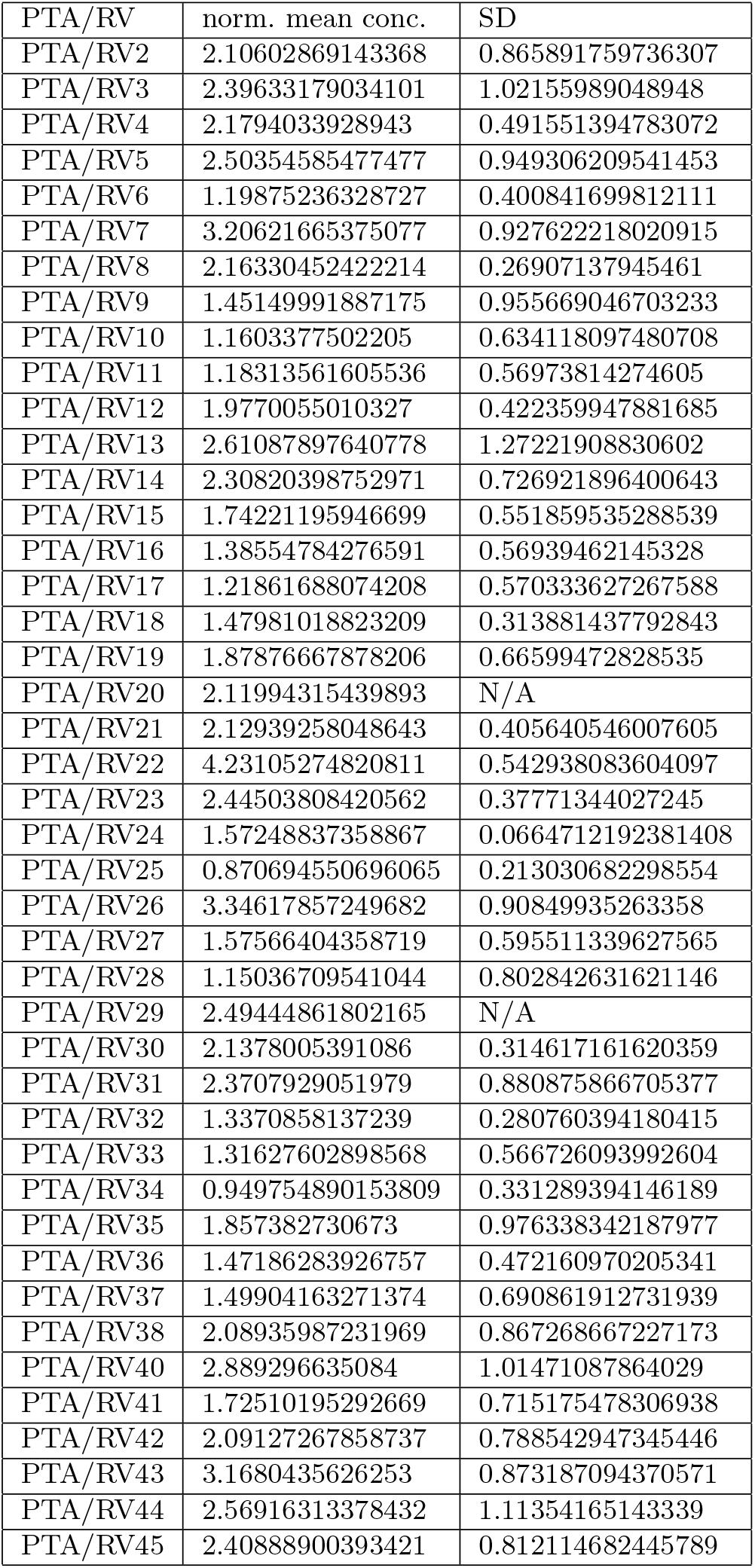
Predicted concentration in PTA/RVs at the endpoint of a 48h E12 kidney organ culture. Simulated normalized mean concentration, 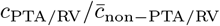, inside each segmented PTA/RV and standard deviation. Data is used for Fig.2F.

### Supplementary Figure Legends

**Figure S1: The geometry effect is largely robust to non-uniform *Wnt9b* expression**. To assess the impact of non-uniform *Wnt9b* expression in the ureteric bud, we solved the steady-state diffusion model on the segmented geometries obtained from LSM, using different *Wnt9b* expression patterns (pink) on the segmented UB geometries, i.e. uniform expression (A), exclusion from tips (B), spotty expression (C), restriction to the cortical side (D) or to the stalks (E). As in Fig.1E, 2C, we use Neumann boundary conditions to model WNT9b influx into the domain. The flux is non-zero in the pink part and zero in the gray parts. The red-white-blue colourmap shows the normalised, integrated WNT9b concentration profile along the normal direction of the UB towards the mesenchyme projected back on the surface of the UB. The surface of segmented PTA/RVs is shown in yellow. (F) Relative predicted WNT9b concentration in segmented PTA/RVs and outside. We obtain similar distributions of relative WT9b up-concentration levels inside the PTA/RVs for all expression patterns, unless *Wnt9b* expression is absent from the corner region facing the PTA/RVs. All simulations were run with a gradient length of λ = 30µm.

**Figure S2: 2D segmentation can result in false impression of branch points**. (A) Widefield kidney (E12) live imaging; transgenic labeling of the ureteric bud (UB) in green, overlaid with the 2D segmentation of the UB used in dynamic simulations (white outline) (Fig.2B). Black arrowheads mark points of under-segmented branches due to the 3D geometry of the tissue. (B) 3D rendered light-sheet data of the UB at the culture endpoint. White arrowheads highlight UB branches that grow below the ureter and are therefore not visible in the 2D segmentation. Scale bars 100µm.

**Figure S3: Comparison of ectopic PTA/RV induced by the artificial zero-flux boundary of the tissue-air interface and by WNT9b-loaded beads**. (A-C) Endpoint of an E11.5 kidney explant (Fig. 3K), cultured for 63 hours in the presence of a WNT9b-soaked bead (marked by white dashed circle in A,C). PTA/RVs were revealed with anti-LHX1 antibodies (yellow), progenitors with anti-SIX2 antibodies (orange), and the UB (green) via the HoxB7-Venus transgene. Ectopic PTA/RVs induced by the artificial zero-flux boundary of the tissue-air interface (A) and by the WNT9b-soaked bead (C) are marked by coloured boxes that correspond to the colour code used in panels D–H. Steady-state diffusion (λ = 30 µm) with uniform secretion of WNT9b from the segmented boundary of the UB (Neumann boundary condition) was solved (B) to evaluate the predicted WNT9b concentration at the position of the detected ectopic PTA/RVs (D–H). Simulated WNT9b concentrations are represented using a colour scale ranging from blue to red. (D–H) Side and front views of the parts boxed in panels A and C, with coloured dashed lines indicating the sectioning. The colour marking of the SIX2-stained volumes vary between orange and brown, depending on the colours that represent other components. PTA/RVs (yellow) are also marked by asterisks. (D–F) The culture-induced PTA/RVs (yellow, marked by asterisks) primarily form near the tissue-air interface on top of the UB (green), where high predicted WNT9b concentrations (red) coincide with SIX2+ NPCs (orange/brown). (G,H) The bead-induced PTA/RVs (yellow, marked by asterisks) develop further away from the UB (green) in regions with predicted lower WNT9b concentrations (blue). (I) Volumes of the PTA/RVs in the end point culture. The timepoint of formation (late to early) was determined using the live imaging data from the culture. The bead-induced PTA/RVs (magenta) exhibited similar volumes and formed at comparable time points as early PTA/RVs (purple). In contrast, the culture-induced PTA/RVs (ciel) had smaller volumes, indicating their emergence at later culture stages.

### Supplementary Movies

**Movie S1:**

**3D geometry of an E12.5 metanephric kidney**. Movie of 3D lightsheet microscopy of an E12.5 kidney and subsequent surface segmentation, which are the basis for downstream analysis. Green is the UB, labelled by HoxB7/myr-Venus expression, orange is SIX2. LHX1, the PTA/RV marker is depicted in yellow.

**Movie S2:**

**3D diffusion simulation on realistic kidney geometry**. 3D diffusion simulation based on image-segmented surfaces. Red areas have high, white medium, and blue low simulated WNT9b concentration. Orange are SIX2-positive cells, yellow are identified PTA/RV. Grey is the kidney cortex. First the simulation is shown in 3D, then it slices through the volume.

**Movie S3:**

**2D kidney explant culture shows the sequential formation of PTA/RV** E12.5 mouse kidneys were cultured for 48h. One frame corresponds to 1h in culture. These images are the basis for the shapes used for the 2D diffusion simulations. Frames were used for Figure 2A.

**Movie S4:**

**Temporal coordination of nephrogenesis and UB branching**. 2D diffusion simulations on segmented live imaging data of kidney explant cultures. Red areas have high, white medium, and blue low simulated WNT9b concentration. Imaging time points where captured once per hour. The growth in-between time points is reconstructed by computing vector fields describing the change in geometry and doing simulations on a continuous growing domain.

**Movie S5:**

**Explant cultures with local WNT9b source**. Time-lapse movies of E11.5 kidneys cultured for 63h. Left: Kidney co-cultured with a control PBS-soaked bead; Middle and right: Kidneys co-cultured with WNT9b-soaked beads. HoxB7/myr-Venus expression is shown in green, brightfield in grey.

### Supplementary Files

**HTML S1:**

**3D diffusion simulation on realistic kidney geometry - interactive plot**. Interactive 3D plot of the simulated WNT9b concentration projected on the UB surface of a E12.5 mouse kidney. Red areas have high, white medium, and blue low simulated WNT9b concentration. Yellow surfaces are PTA/RV.

**HTML S2:**

**3D simulation on cultured kidney geometry**. Interactive 3D plot of the simulated WNT9b concentration projected on the UB surface of a explant culture endpoint after 48h. Red areas have high, white medium, and blue low simulated WNT9b concentration. Yellow surfaces are PTA/RV.

