## Supplementary Text & Figures for "Geometric Effects Position Renal Vesicles During Kidney Development"

### Supplemental material

#### Supplementary Data

**Table S1: Predicted concentration in the PTA/RVs of an E12.5 kidney.** Simulated normalized mean concentration, $c_{PTA/RV}/\bar{c}_{SIX2}$, inside each segmented PTA/RV and standard deviation. From the E12.5 Kidney used for Fig.1D.

| PTA/RV | norm. mean conc. | SD |
|---|---|---|
| PTA/RV1 | 2.10460123167059 | 0.784301184498161 |
| PTA/RV2 | 1.72093520735421 | 0.462866825453608 |
| PTA/RV3 | 2.28654758432875 | 0.816871034942931 |
| PTA/RV4 | 1.98017417670302 | 0.83720111501182 |
| PTA/RV5 | 2.23117441526323 | 0.689618259265129 |
| PTA/RV6 | 2.3568583628366 | 0.523467432081343 |
| PTA/RV7 | 1.98283250798053 | 0.82473092806899 |
| PTA/RV8 | 1.84812853602427 | 0.953668436073596 |
| PTA/RV9 | 1.57182549499172 | 0.872044894290876 |
| PTA/RV10 | 1.9628332678972 | 0.75035306715877 |
| PTA/RV11 | 2.39424794830583 | 0.549865863560481 |
| PTA/RV12 | 1.93922717479229 | 0.738153807556894 |
| PTA/RV13 | 2.1449830693159 | 0.800876969300404 |
| PTA/RV14 | 2.20504185023655 | 0.767819761191748 |

**Table S2: Predicted concentration in PTA/RVs at the endpoint of a 48h E12 kidney organ culture.** Simulated normalized mean concentration, $c_{\mathrm{PTA/RV}}/\bar{c}_{\mathrm{non-PTA/RV}}$, inside each segmented PTA/RV and standard deviation. Data is used for Fig.2F.

| PTA/RV | norm. mean conc. | SD |
|---|---|---|
| PTA/RV2 | 2.10602869143368 | 0.865891759736307 |
| PTA/RV3 | 2.39633179034101 | 1.02155989048948 |
| PTA/RV4 | 2.1794033928943 | 0.491551394783072 |
| PTA/RV5 | 2.50354585477477 | 0.949306209541453 |
| PTA/RV6 | 1.19875236328727 | 0.400841699812111 |
| PTA/RV7 | 3.20621665375077 | 0.927622218020915 |
| PTA/RV8 | 2.16330452422214 | 0.26907137945461 |
| PTA/RV9 | 1.45149991887175 | 0.955669046703233 |
| PTA/RV10 | 1.1603377502205 | 0.634118097480708 |
| PTA/RV11 | 1.18313561605536 | 0.56973814274605 |
| PTA/RV12 | 1.9770055010327 | 0.422359947881685 |
| PTA/RV13 | 2.61087897640778 | 1.27221908830602 |
| PTA/RV14 | 2.30820398752971 | 0.726921896400643 |
| PTA/RV15 | 1.74221195946699 | 0.551859535288539 |
| PTA/RV16 | 1.38554784276591 | 0.56939462145328 |
| PTA/RV17 | 1.21861688074208 | 0.570333627267588 |
| PTA/RV18 | 1.47981018823209 | 0.313881437792843 |
| PTA/RV19 | 1.87876667878206 | 0.66599472828535 |
| PTA/RV20 | 2.11994315439893 | N/A |
| PTA/RV21 | 2.12939258048643 | 0.405640546007605 |
| PTA/RV22 | 4.23105274820811 | 0.542938083604097 |
| PTA/RV23 | 2.44503808420562 | 0.37771344027245 |
| PTA/RV24 | 1.57248837358867 | 0.0664712192381408 |
| PTA/RV25 | 0.870694550696065 | 0.213030682298554 |
| PTA/RV26 | 3.34617857249682 | 0.90849935263358 |
| PTA/RV27 | 1.57566404358719 | 0.595511339627565 |
| PTA/RV28 | 1.15036709541044 | 0.802842631621146 |
| PTA/RV29 | 2.49444861802165 | N/A |
| PTA/RV30 | 2.1378005391086 | 0.314617161620359 |
| PTA/RV31 | 2.3707929051979 | 0.880875866705377 |
| PTA/RV32 | 1.3370858137239 | 0.280760394180415 |
| PTA/RV33 | 1.31627602898568 | 0.566726093992604 |
| PTA/RV34 | 0.949754890153809 | 0.331289394146189 |
| PTA/RV35 | 1.857382730673 | 0.976338342187977 |
| PTA/RV36 | 1.47186283926757 | 0.472160970205341 |
| PTA/RV37 | 1.49904163271374 | 0.690861912731939 |
| PTA/RV38 | 2.08935987231969 | 0.867268667227173 |
| PTA/RV40 | 2.889296635084 | 1.01471087864029 |
| PTA/RV41 | 1.72510195292669 | 0.715175478306938 |
| PTA/RV42 | 2.09127267858737 | 0.788542947345446 |
| PTA/RV43 | 3.1680435626253 | 0.873187094370571 |
| PTA/RV44 | 2.56916313378432 | 1.11354165143339 |
| PTA/RV45 | 2.40888900393421 | 0.812114682445789 |

#### Supplementary Figures

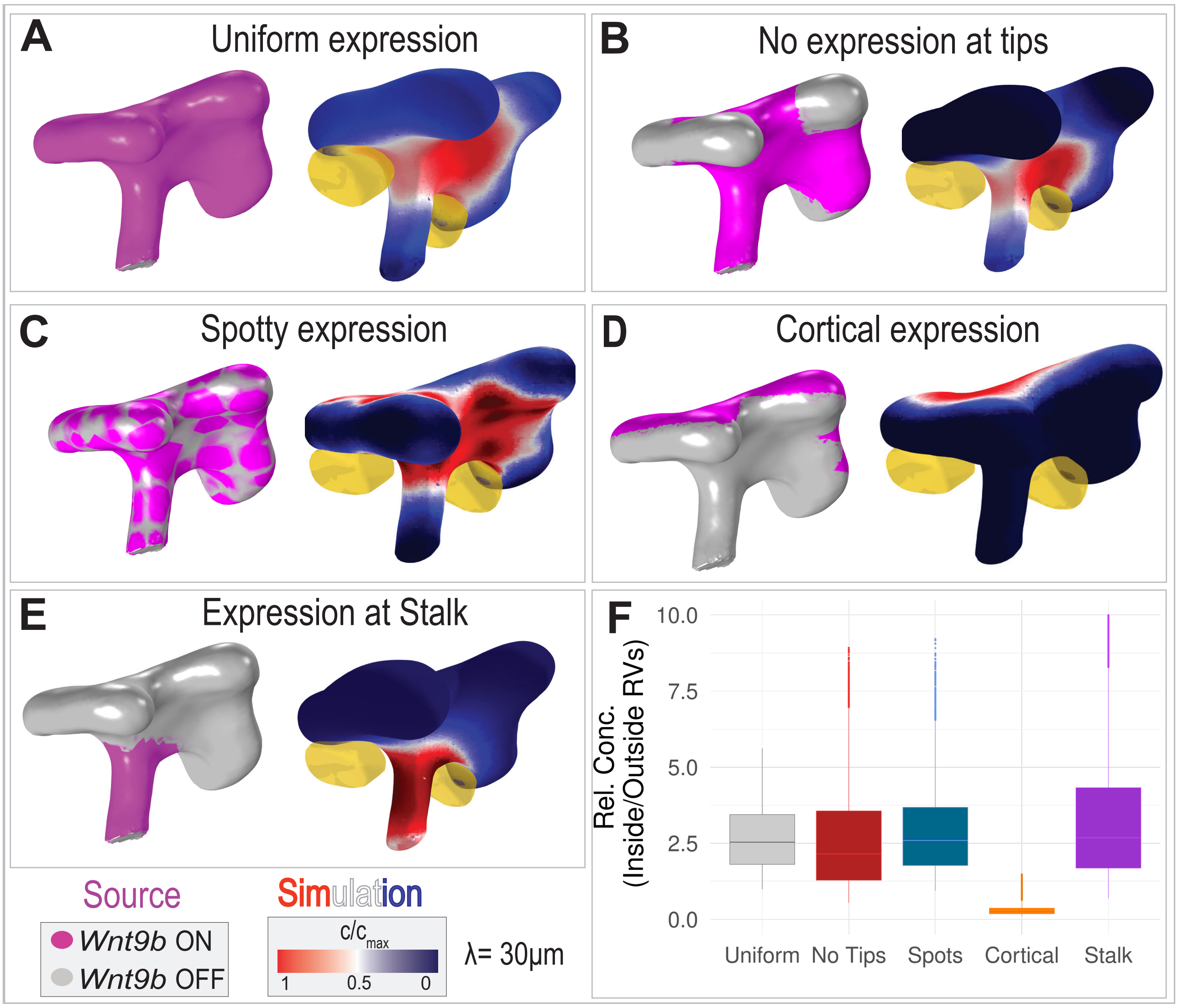


**Figure S1: The geometry effect is largely robust to non-uniform *Wnt9b* expression.** To assess the impact of non-uniform *Wnt9b* expression in the ureteric bud, we solved the steady-state diffusion model on the segmented geometries obtained from LSM, using different *Wnt9b* expression patterns (pink) on the segmented UB geometries, i.e. uniform expression (A), exclusion from tips (B), spotty expression (C), restriction to the cortical side (D) or to the stalks (E). As in Fig.1E, 2C, we use Neumann boundary conditions to model WNT9b influx into the domain. The flux is non-zero in the pink part and zero in the gray parts. The red-white-blue colourmap shows the normalised, integrated WNT9b concentration profile along the normal direction of the UB towards the mesenchyme projected back on the surface of the UB. The surface of segmented PTA/RVs is shown in yellow. (F) Relative predicted WNT9b concentration in segmented PTA/RVs and outside. We obtain similar distributions of relative WT9b up-concentration levels inside the PTA/RVs for all expression patterns, unless *Wnt9b* expression is absent from the corner region facing the PTA/RVs. All simulations were run with a gradient length of $\lambda = 30\mu m$.

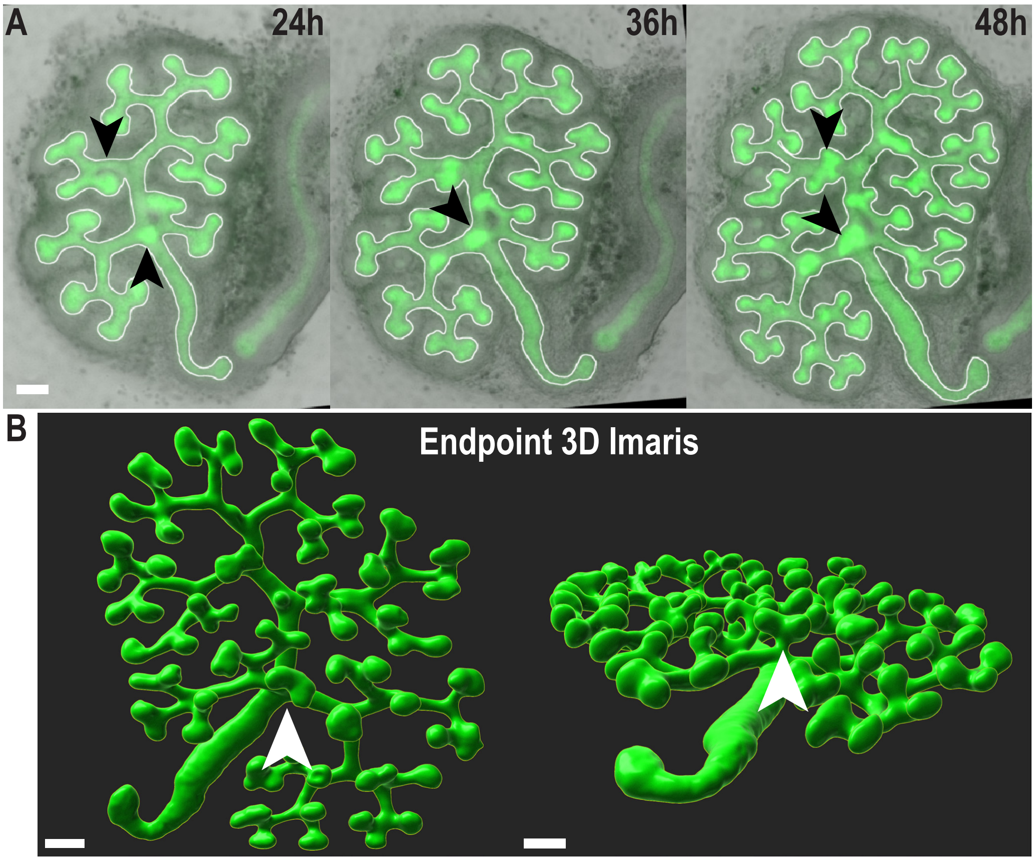


**Figure S2: 2D segmentation can result in false impression of branch points.** (A) Widefield kidney (E12) live imaging; transgenic labeling of the ureteric bud (UB) in green, overlaid with the 2D segmentation of the UB used in dynamic simulations (white outline) (Fig.2B). Black arrowheads mark points of under-segmented branches due to the 3D geometry of the tissue. (B) 3D rendered light-sheet data of the UB at the culture endpoint. White arrowheads highlight UB branches that grow below the ureter and are therefore not visible in the 2D segmentation. Scale bars 100µm.

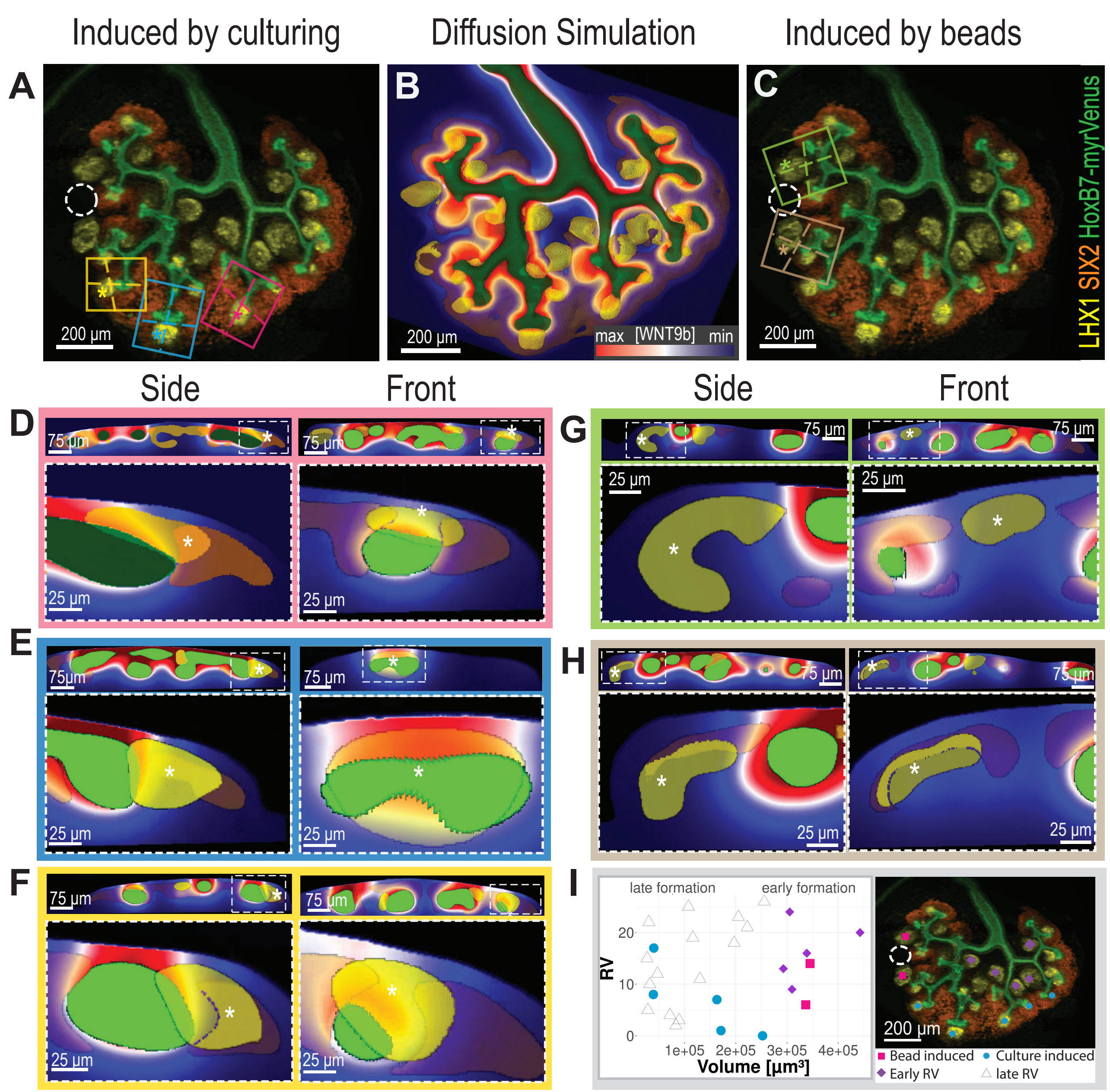


**Figure S3: Comparison of ectopic PTA/RV induced by the artificial zero-flux boundary of the tissue-air interface and by WNT9b-loaded beads.** (A-C) Endpoint of an E11.5 kidney explant (Fig. 3K), cultured for 63 hours in the presence of a WNT9b-soaked bead (marked by white dashed circle in A,C). PTA/RVs were revealed with anti-LHX1 antibodies (yellow), progenitors with anti-SIX2 antibodies (orange), and the UB (green) via the HoxB7-Venus transgene. Ectopic PTA/RVs induced by the artificial zero-flux boundary of the tissue-air interface (A) and by the WNT9b-soaked bead (C) are marked by coloured boxes that correspond to the colour code used in panels D–H. Steady-state diffusion ($\lambda = 30$ µm) with uniform secretion of WNT9b from the segmented boundary of the UB (Neumann boundary condition) was solved (B) to evaluate the predicted WNT9b concentration at the position of the detected ectopic PTA/RVs (D–H). Simulated WNT9b concentrations are represented using a colour scale ranging from blue to red. (D–H) Side and front views of the parts boxed in panels A and C, with coloured dashed lines indicating the sectioning. The colour marking of the SIX2-stained volumes vary between orange and brown, depending on the colours that represent other components. PTA/RVs (yellow) are also marked by asterisks. (D–F) The culture-induced PTA/RVs (yellow, marked by asterisks) primarily form near the tissue-air interface on top of the UB (green), where high predicted WNT9b concentrations (red) coincide with SIX2+ NPCs (orange/brown). (G,H) The bead-induced PTA/RVs (yellow, marked by asterisks) develop further away from the UB (green) in regions with predicted lower WNT9b concentrations (blue). (I) Volumes of the PTA/RVs in the end point culture. The timepoint of formation (late to early) was determined using the live imaging data from the culture. The bead-induced PTA/RVs (magenta) exhibited similar volumes and formed at comparable time points as early PTA/RVs (purple). In contrast, the culture-induced PTA/RVs (ciel) had smaller volumes, indicating their emergence at later culture stages.
